# Optic Nerve Tortuosity and Globe Proptosis in Normal and Glaucoma Subjects

**DOI:** 10.1101/397570

**Authors:** Xiaofei Wang, Helmut Rumpel, Mani Baskaran, Tin A Tun, Nicholas Strouthidis, Shamira A Perera, Monisha E Nongpiur, Winston Eng Hoe Lim, Tin Aung, Dan Milea, Michaël J. A. Girard

## Abstract

**Purpose:** To assess the difference in optic nerve tortuosity during eye movements and globe proptosis between primary open angle glaucoma and normal subjects using orbital magnetic resonance imaging.

**Methods:** 10 Chinese subjects matched for age, ethnicity and refractive errors were recruited, including five normal controls and five patients with primary open angle glaucoma. All subjects underwent magnetic resonance imaging to assess their optic nerves and globes for three eye positions: primary gaze, adduction and abduction. Optic nerve tortuosity (optic nerve length divided by the distance between two ends) and globe proptosis (maximum distance between cornea and interzygomatic line) were measured from magnetic resonance imaging images.

**Results:** In adduction, the tortuosity of normal eyes was significantly larger than that of the glaucomatous eyes. Optic nerve tortuosity in adduction in the control and glaucoma groups were 1.004±0.003 (mean ± standard deviation) and 1.001±0.001, respectively (p=0.037). Globe proptosis (primary gaze) in glaucoma subjects (19.14±2.11 mm) was significantly higher than that in control subjects (15.32±2.79 mm; p = 0.046).

**Conclusions:** In this sample, subjects with glaucoma exhibited more taut optic nerves and more protruding eye globes compared to normal eyes. This may impact optic nerve head deformations in anatomically predisposed patients.

**Précis:** Eyes with glaucoma have tauter optic nerves compared with normal eyes, which may exert more force on the optic nerve head tissues during eye movements.

## INTRODUCTION

The main site of retinal ganglion cell (RGC) damage in glaucoma is the optic nerve head (ONH), especially the lamina cribrosa (LC) within the ONH. To date, the pathogenesis of glaucoma is not fully elucidated. We know that elevated intraocular pressure (IOP) is the major risk factor for glaucoma.^1, 2^ However, glaucoma also occurs in patients with IOP within normal levels,^3^ and may not develop in those with elevated IOP. In short, our understanding of glaucoma is insufficient: While IOP is crucial, it is likely that risk factors other than IOP play an important role in the development and progression of this blinding disease.

The biomechanical theory of glaucoma hypothesizes that elevated IOP deforms the ONH tissues, and that these deformations (directly or indirectly) drive RGC death.^4^ This theory is supported by much circumstantial evidence^5-7^, and can explain differing sensitivities to IOP. However, IOP is not the only load that can induce ONH deformations. For instance, the cerebrospinal fluid pressure (CSFP) in the optic nerve (ON) subarachnoid space can deform the ONH, and in fact, low CSFP is thought to be linked with the development of normal tension glaucoma.^8^ Recently, eye movements have also been shown to significantly deform the ON and ONH tissues. Specifically, recent studies that employed optical coherence tomography (OCT),^9-12^ computational modeling,^13-15^ and magnetic resonance imaging (MRI) ^16, 17^ all converge to the finding that the optic nerve (ON) can pull the back of the eye during eye movements to result in large deformations within the optic nerve head (ONH) tissues. ^13, 15, 16^

There is now a growing interest to understand whether ONH deformations induced by the ON traction force could initiate the development and progression of optic neuropathies including glaucoma.^9, 11, 12, 16, 18^ Using FE modeling, we have previously predicted that a stiff dura would generate large ONH deformations.^13^ Moreover, and from a strict mechanical point of view, we have hypothesized that a smaller amount of ON slack in primary gaze (i.e. a less tortuous ON) would intuitively result in larger ONH deformations during eye movements due to a rapid straightening and pulling effect of the ON. While it is not yet feasible to measure the stiffness of the dura in vivo, ON tortuosity can be assessed with MRI. A recent study has reported that optic nerves of normal tension glaucoma patients are more tortuous than those of normal controls.^17^ However, no studies have investigated the optic nerve morphology in high-tension glaucoma in Chinese subjects.

The aim of this study was to measure ON tortuosity and globe proptosis in healthy and subjects with primary open glaucoma (POAG) in different gaze positions using MRI.

## METHODS

### Subject Recruitment & Clinical Tests

For this study, ten Chinese subjects were recruited from the Singapore National Eye Center. Among these, five were normal controls and five were diagnosed with primary open angle glaucoma (POAG); 3 were female and 7 were male. Normal controls were recruited from Singapore National Eye Center staff and the general public by advertising. Inclusion criteria for normal subjects were: age ≥ 50 years, phakic, and with refractive errors within the range of ±3.5 Diopters. POAG cases were diagnosed by the presence of glaucomatous optic neuropathy, defined as vertical cup-to-disc ratio of >0.7 and/or neuroretinal rim narrowing with an associated visual field defect on standard automated perimetry. Visual field loss for POAG cases should less than −6 dB mean deviation value. The glaucoma subjects were newly diagnosed or receiving pharmacologic treatment. We excluded subjects with thyroid eye disease, a history of intraocular surgery, strabismus, ocular motor palsies, orbital or brain tumors, orbital or brain surgeries and subjects that were not suitable for MRI scanning.

All the subjects underwent the following ocular examinations for both eyes: 1) measurement of refraction using an autokeratometer (RK-5, Canon, Tokyo, Japan); 2) measurement of vertical cup-to-disc ratio using a slit lamp. 3) measurement of visual acuity using a logMar chart; 4) anterior segment swept source OCT imaging (SS-1000, CASIA, Tomey, Nagoya, Japan) and 5) measurement of IOP using a Goldmann applanation tonometry (AT900D; Haag-Streit, Köniz, Switzerland). Visual fields were assessed by standard automated perimetry (SITA-FAST 24-2 program; Humphrey Field Analyzer II-750i, Carl Zeiss Meditec, Dublin, CA, USA) for both eyes in glaucoma subjects.

The protocol was approved by the SingHealth Centralized Institutional Review Board and adhered to the ethical principles outlined in the Declaration of Helsinki. Written voluntary informed consent was obtained from each subject.

### Magnetic Resonance Imaging

Each subject’s orbital regions were imaged with a 3T MRI scanner and a 32- channel head coil (Magnetom Skyra, Siemens Healthcare Sector, Erlangen, Germany) without the use of any contrast agent. FL3D (3D flash sequence) used the following parameters: TR 10 ms, TE 3 ms, flip angle 22°, FOV 140 mm, matrix 256 x256 x 8, resulting in the axial images with an in-plane resolution of 0.5 mm per pixel and a slice thickness of 2 mm. With 60% oversampling and two averages, the acquisition time for each scan was 1.06 min. The location of the MRI slice was carefully determined during acquisition so that it divided the ON and the eye globe into superior and inferior halves.

During the MRI procedure, the subjects were orally instructed to direct their gaze on one of the three visual targets, allowing image acquisition in the primary position (baseline), and on left and right gaze (fixed order). The exact eye rotation amplitudes were measured from MRI images in the post-processing stage. These three visual targets were around 32 cm from subjects’ eyes and were 1 inch in width (visual angle of about 4.5°) to achieve a very good visibility. After the subjects were positioned in the MRI scanner, they were trained to direct their gaze on individual targets before the MRI session.

### Image Delineation

For each eye, the obtained 2D MRI image was delineated by a single grader (XW) for further analysis using custom-written software in MATLAB (Version 2015a; Mathworks, Inc., Natick, MA, USA). Six structures were delineated manually (**Figure 1A**): the anterior cornea (in green), the lens capsule and center (yellow), the scleral shell (in red), the temporal and nasal boundaries of the ON (in yellow), and the interzygomatic line (in green). Specifically, for the cornea, ∼6 points were marked that were fitted with a quadratic polynomial. For the lens capsule, ∼10 points were marked that were fitted with an ellipse; the ellipse center was then assumed to be that of the lens. The scleral shell was also identified (∼30 discrete points) and fitted with an ellipse to approximate the globe. For the ON, ∼10 discrete points were marked to identify its temporal or nasal boundary, which were then connected using a cubic spline. Finally, two points at the anterior limits of the left and right zygomatic bones were identified and a line that passed through those 2 points was defined as the interzygomatic line.^19^

A central line passing through the ON was derived from the delineated temporal and nasal ON boundaries, from which we extracted its tortuosity (defined as ON tortuosity).

**Figure 1.**
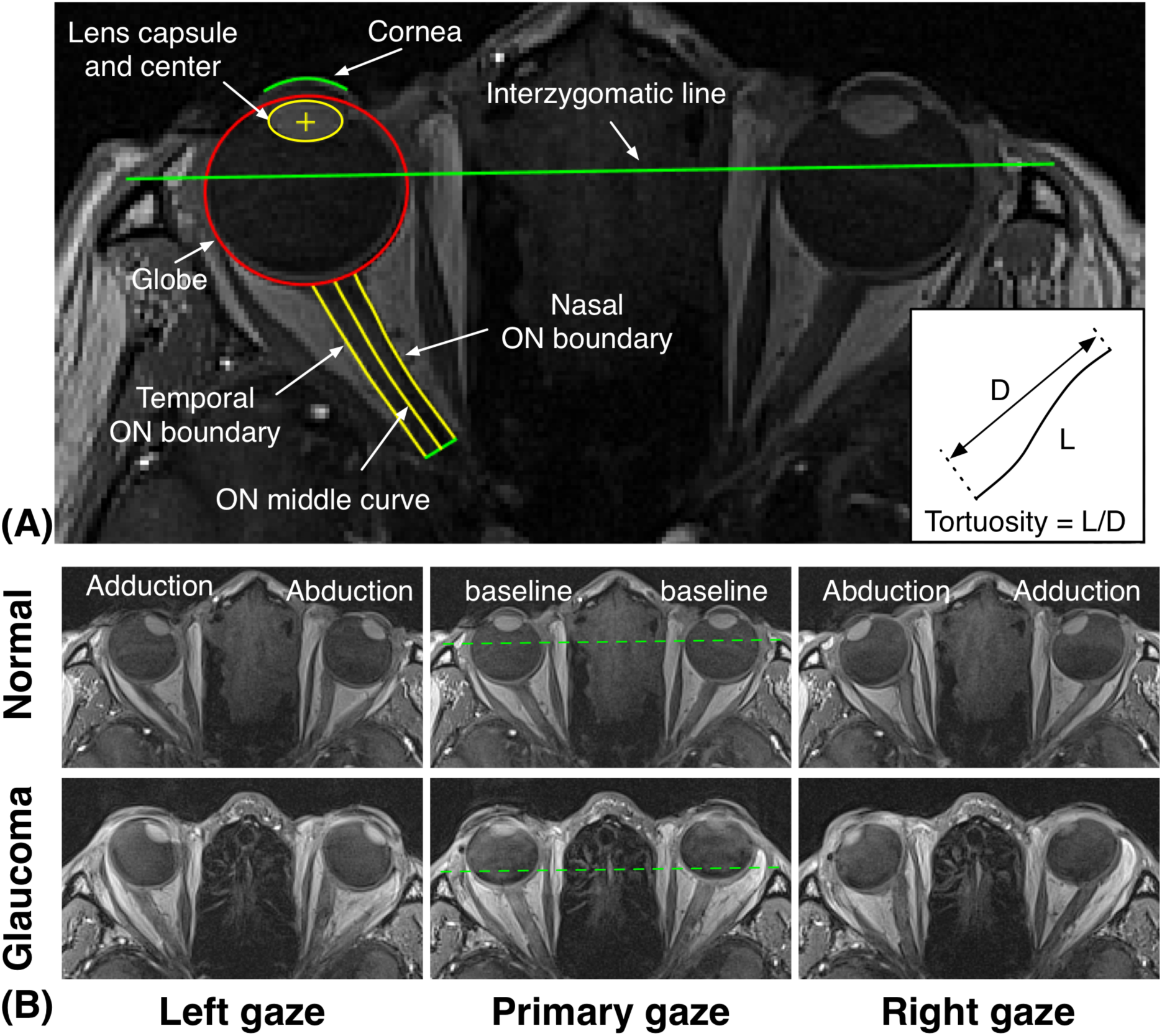
(**A**) Delineated features of the eye and the orbit. All measurements were calculated automatically using our custom-written application based on these features. Figure on the right corner indicates tortuosity definition. (**B**) Axial MRI images for a glaucoma and a normal subject in the primary, left and right gaze positions. Green dashed lines indicate interzygomatic lines. Note the high proptosis in the glaucoma subject.

### Measurement of Optic Nerve Tortuosity

The amount of ON ‘slack’ is an important factor that may influence the magnitude of the ON traction force during eye movements. For instance, a smaller amount of slack (i.e. a less tortuous ON) in some individuals may result in larger ONH deformations during eye movements due to a rapid straightening effect of the ON. To this end, we measured the tortuosity of the ON for each eye. As the posterior end of the intraorbital ON were faint, tortuosity was calculated based on a 20 mm long ON segment. ON tortuosity was defined as the length of the ON central line segment (20 mm for all eyes in this study) divided by the distance between two end points of the line segment (minimum path in the axial MRI image). The calculations of length and distance were automatically performed in MATLAB. We reported ON tortuosities for all eyes in all three gaze positions. Note that the ON tortuosity was calculated based on the fitted boundaries of the globe and ON. Therefore, subpixel resolution can be achieved in this measurement.

### Measurement of Globe Proptosis on MRI

We also measured the globe proptosis on axial MRI scans, in baseline gaze position, which was defined as the maximum distance between the cornea and the interzygomatic line. The distance was calculated automatically by our application using the delineated structures (**Figure 1A**).

### Statistical Analysis

As data from both eyes were used, linear mixed models were employed to adjust for inter-eye correlations. All statistical analyses were performed using R (R Foundation, Vienna, Austria).

## RESULTS

### Subjects

The mean age (±SD) for all subjects at the time of MRI scanning was 61.6±7.6 years. The mean IOP for POAG eyes before and after IOP lowering treatment were 26.4±4.6 mmHg and 14.1±2.3 mmHg, respectively. The mean IOP for normal eyes was 15.3±3.6 mmHg. Eye rotation angles were 31±5° for abduction and 37±4° for abduction as measured from the MRI images.

### Axial MRI Images

**Figure 1B** shows axial orbital MRI images in a glaucoma patient and in a healthy control, fixating the central, the left and the right visual targets. In adduction, the ONs were highly stretched and were tauter than in normal positions. In abduction, the ONs were also stretched but were more tortuous than those in adduction.

### Optic Nerve Tortuosity

#### Normal vs Glaucoma

ON tortuosity in baseline gaze, adduction and abduction of 30° are summarized in **Figure 2** as box plots for both control and glaucoma groups (note that the ON central line was also drawn for each subject to ease interpretation of our results). In each position, the ON tortuosity of normal eyes was significantly larger than that of the glaucomatous eyes. In baseline gaze, ON tortuosity (mean±SD) in the control and glaucoma groups was 1.013±0.019 and 1.002±0.003, respectively (p=0.248); in adduction, 1.004±0.003 and 1.001±0.001, respectively (p=0.037); in abduction, 1.033±0.034 and 1.011±0.009, respectively (p=0.215).

**Figure 2.**
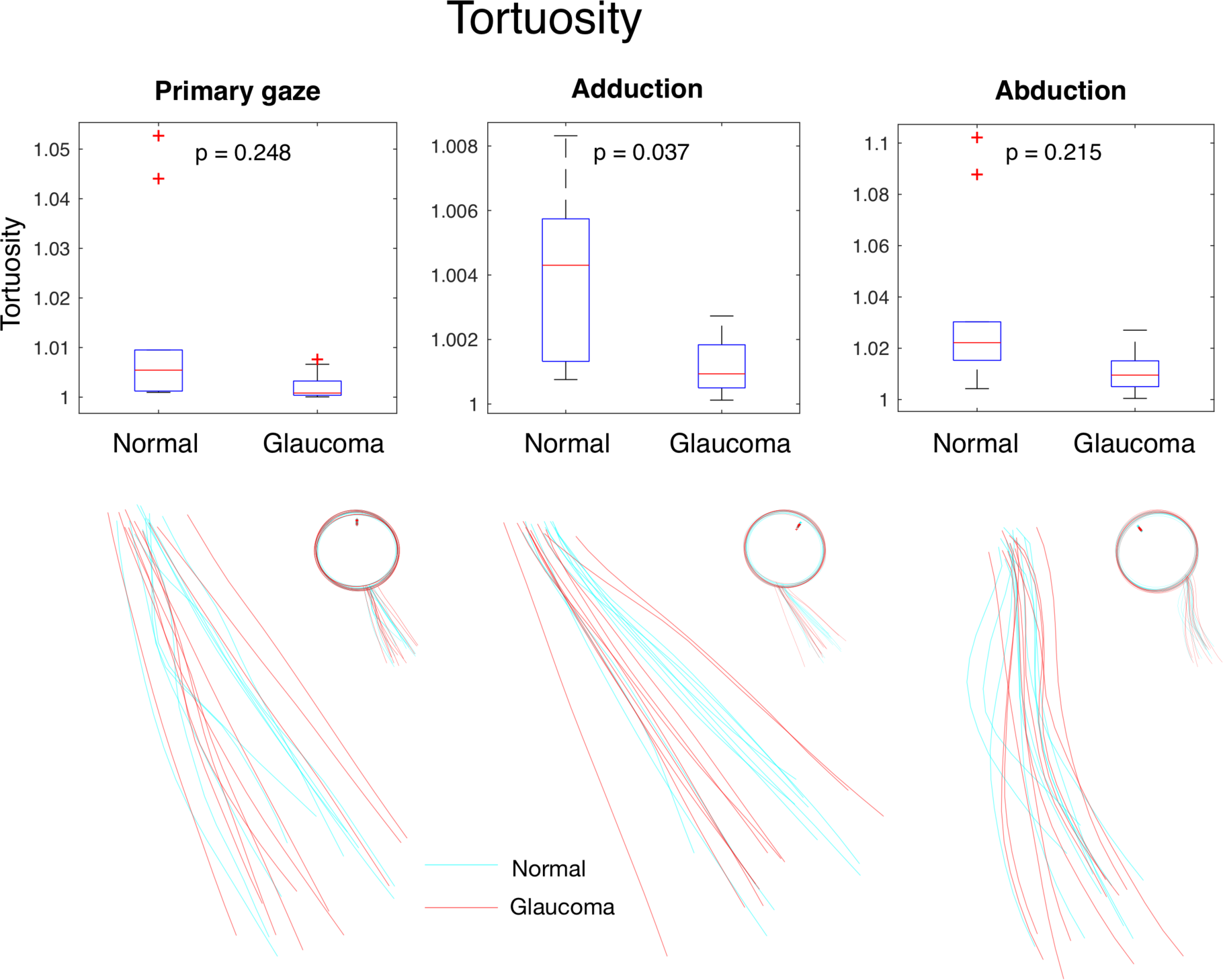
**Upper row**: box plots of ON tortuosities for normal and glaucoma subjects in primary gaze, adduction and abduction. ON tortuosities of normal eyes in adduction were statistically higher than those of glaucoma subjects. **Lower row**: delineated ON midlines at various eye positions.

#### Adduction vs Abduction vs Baseline Gaze

For all eyes, ON tortuosity in adduction (1.003±0.003) was on average smaller than that in baseline gaze (1.008±0.014; p=0.0423). In abduction, ON tortuosity (1.022±0.027) was significantly larger than that in baseline (p<0.001) and in adduction (p<0.001).

### Globe Proptosis

#### Proptosis

We found that globe proptosis (baseline gaze) in glaucoma subjects (19.14±2.11 mm) was significantly higher than that in control eyes (15.32±2.79 mm; p = 0.046; **Figure 3**).

**Figure 3.**
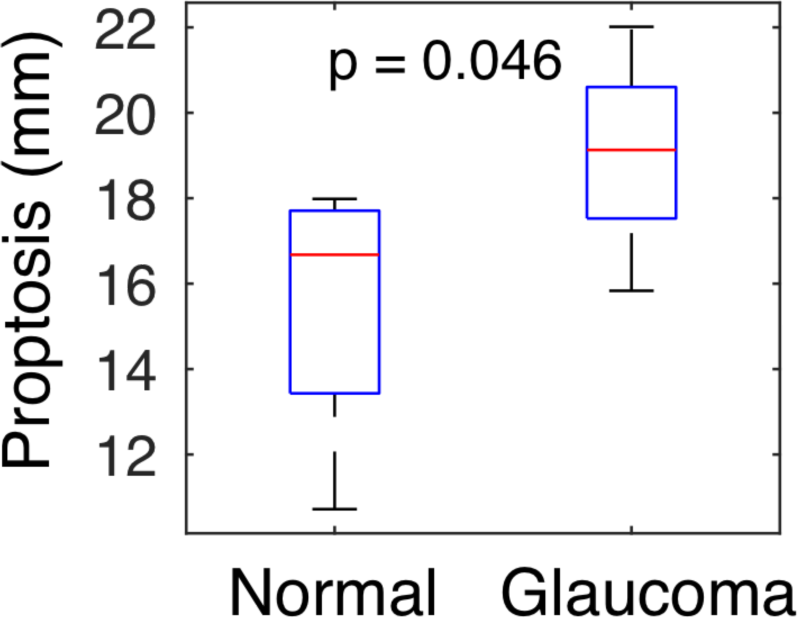
Box plots of globe proptosis for normal and glaucoma subjects in primary gaze. Proptoses of glaucomatous eyes were higher than those of the normal eyes.

## DISCUSSION

Glaucoma is a heterogeneous disease with a pathogenesis that may involve many factors such as age, ethnicity, genes, IOP and cerebrospinal fluid pressure (CSFP) ^20^. ONH deformations induced by IOP or CSFP and their correlations with glaucoma have been studied extensively ^21^. Eye movements can also deform the ONH significantly ^9, 11-13, 16^, but to date, a potential biomechanical link with glaucoma remains unclear. A recent study ^17^has reported that the ON in patients with normal tension glaucoma are more tortuous than those of controls. However, no studies have investigated the optic nerve morphology in high-tension POAG in Chinese subjects. Our study suggests that ONs in glaucoma subjects are more taut than ONs in normal controls in adduction. The implications of such a finding are still unclear, but a more taut ON has the potential to result in an earlier extinction of ON redundancy during eye movements and cause greater stretch within the ON and ONH. However, ONH deformation levels are also dependent on the biomechanical properties of the dura mater and of the peripapillary sclera ^13, 15^. Whether a more taut ON would be associated with higher glaucoma susceptibility still remains speculative.

In this study, we found that globe proptosis in normal subjects was 15.32±2.79 mm, which is highly consistent with data obtained from a large cohort of Chinese adults^22^(15.3±2.3 mm for patients aged between 70 and 87 years old; 15.7±1.8 for patients aged between 20 and 69 years old; measured with Hertel Exophthalmometry^23^). Interestingly, we also found that globe proptosis in glaucoma subjects (19.14±2.11 mm; **Figure 3**) was conspicuously higher than in normal eyes. This difference in proptosis was not caused by elongation of the globe as we found no significant difference in axial length between normal and glaucoma groups (**Supplementary material A**). We believe this is an interesting but controversial result that would need to be validated in a larger population.

In adduction, ON tortuosity was smaller than that in baseline gaze. This finding is intuitive as the ON is highly stretched in adduction and thus less curved. On the contrary, ON tortuosity in abduction for both glaucoma and normal groups was larger than that in baseline gaze. This result may appear counterintuitive because, in abduction, the ON is likely to exhibit a certain level of stretch and the distance between the ONH and the orbital apex is increasing.^12^ This phenomenon could potentially be explained by the combination of: (1) a smaller ON traction force in abduction (as predicted through finite element modeling^15^); and (2) resistance from the orbital tissues (fat and extraocular muscles) to restrict the ‘free movement’ of the ON in abduction. However, to date, this resistance effect has not been well studied. It is possibly very complex due to nonlinear viscoelastic material properties of the orbital fat, and heterogeneous frictional effects between the orbital fat and the wall of the ON.^24^ Several research groups have suggested that the orbital fat, although soft, is able to provide essential supports for stabilizing the extraocular muscle bellies during eye movements,^25, 26^ while others have suggested that the orbital fat is more like a fluid with no resistance.^16^ Additionally, changes in extraocular muscle shapes and volumes (induced by contraction and relaxation) during eye movements may influence the deformations of surrounding tissues and may subsequently affect ON deformations. Further studies are needed to investigate the biomechanical properties of the human orbital fat and their interactions with the ON and extraocular muscles.

The prevalence of high tension glaucoma, normal tension glaucoma and ocular hypertension has been reported to be significantly higher in patients with thyroid eye disease (TED; which may cause globe protrusion) than in the general population.^27-30^ The exact mechanisms of ocular hypertension in TED are not completely understood and were hypothesized to be caused by an increase in episcleral venous pressure induced by elevated retrobulbar pressure^31^and mucopolysaccharide deposition in the trabecular meshwork.^29^However, why a higher prevalence of normal tension glaucoma is observed in TED cannot yet be explained. Other factors, other than IOP, may be involved. We speculate that prolonged proptosis and/or muscle enlargement could restrict the free motion of the ON during eye movements, which may in turn result in more tension transmitted to the ONH and induce axonal insult. We also believe that the effects of IOP, CSFP, or optic nerve traction are not necessarily exclusive but complementary. For instance, gaze-induced ONH deformations in normal tension glaucoma could play a larger role than IOP while in high tension glaucoma the effects of IOP is more prominent. Moreover, the interplay of ONH loads (i.e. IOP, CSFP and ON traction) may also be of importance in our understanding of glaucoma mechanisms. For instance, Sibony has demonstrated that elevated CSFP in papilledema can increase gaze-induced ONH deformations.^11^ Furthermore, it is also possible that prolonged abnormal stretches of the globe by the ON during eye movements may alter IOP but this has not yet been fully explored, other than in monkeys.^32^ Finally, IOP and CSFP are chronic loads while eye movements are transient in nature, thus the consequences of high ON traction is likely to be different. Our eyes exhibit frequent movements in daily activities (around 170,000 saccades per day^33^) and during sleep. The links between frequent transient ONH deformation during eye movements and corresponding tissue damage and remodeling are yet to be established.

***Limitations.*** In this study, several limitations warrant further discussion. First, only 10 subjects were recruited. While the sample size is small, all patients were matched for age, ethnicity, and refractive errors to limit the influence of these co-founding factors.

Second, only one 2D axial image was analyzed, and we were not able to capture the potential curvatures of the ON in the superior-inferior direction. However, our preliminary study using 20 young subjects (unpublished data) showed that horizontal eye movement could only cause changes of ON tortuosity in the horizontal plane. Therefore, a 2D MRI image is capable to capture the feature of ON tortuosity during horizontal eye movement. Further studies may use high spatial resolution images to study the 3D shape changes of ON during eye movements.

Third, the length of the ON was not measured in this study because we were not able to identify the optic canal precisely from our MRI scans. Although an estimate is possible using the medial rectus muscle end, the values may not be reliable. The intraorbital ON length may be a very important factor that influences ON stretching during eye movement. Future studies should include this important parameter whenever possible.

## CONCLUSIONS

In conclusion, we demonstrated a more taut ON and a more protruding globe in glaucoma subjects in a small population, which may result in larger deformations of the ONH during eye movements.

## Supplementary material A

Axial lengths were calculated based on delineated features of the globe. Specifically, an axial line was drawn through the globe center and the lens center. The distance between the intersections of the axial line with the cornea and the posterior globe was defined as the axial length. The measured axial length for glaucoma and normal eyes was 25.68±1.11 and 25.13±0.77 mm, respectively. There was no statistical difference in axial length between normal and glaucomatous eyes (p = 0.31).

